# Amphotericin B resistance in *Leishmania mexicana*: Alterations to sterol metabolism, lipid transport and oxidative stress response

**DOI:** 10.1101/2021.12.08.471712

**Authors:** Edubiel A. Alpizar-Sosa, Nur Raihana Binti Ithnin, Wenbin Wei, Andrew W. Pountain, Stefan K. Weidt, Anne M. Donachie, Ryan Ritchie, Emily A. Dickie, Richard J. S. Burchmore, Paul W. Denny, Michael P. Barrett

## Abstract

Amphotericin B is increasingly used in treatment of leishmaniasis. Here, fourteen independent lines of *Leishmania mexicana* and one *L. infantum* line were selected for resistance to either amphotericin B or the related polyene antimicrobial, nystatin. Sterol profiling revealed that, in each line, the predominant ergostane-type sterol of wild-type cells was replaced by other sterol species. Broadly, two different profiles emerged among the resistant lines. Whole genome sequencing then showed that these distinct profiles were due either to mutations in the sterol methyl transferase (C24SMT) gene locus or the sterol C5 desaturase (C5DS) gene. In three lines an additional deletion of the miltefosine transporter was found. Differences in sensitivity to amphotericin B were apparent, depending on whether cells were grown in HOMEM, supplemented with foetal bovine serum, or a serum free defined medium (DM). These differences appeared to relate to the presence of lipids in the former. Metabolomic analysis after exposure to AmB showed that a large increase in glucose flux via the pentose phosphate pathway preceded cell death in cells sustained in HOMEM but not DM, indicating the oxidative stress was more significantly induced under HOMEM conditions. Several of the lines were tested for ability to infect macrophages and replicate as amastigote forms, alongside their ability to establish infections in mice. While several lines showed reduced virulence, at least one AmB resistant line displayed heightened virulence in mice whilst retaining its resistance phenotype, emphasising the risks of resistance emerging to this critical drug.

## Introduction

The Leishmaniases are caused by parasitic protozoa of the genus *Leishmania* and are transmitted via the bite of infected female phlebotomine sand flies. Clinical forms range from self-healing but disfiguring cutaneous (CL) and mucocutaneous forms (MCL) (1–3) to potentially fatal visceral leishmaniasis (VL) (2), which can also lead to post-kala-azar dermal leishmaniasis (PKDL) in some instances following treatment (4). No vaccine against human leishmaniasis exists (5), and patient management relies on chemotherapy which is currently unsatisfactory for reasons including cost, toxicity (6, 7), drug resistance (1,8–10) and duration and mode of administration. Amphotericin B (AmB) is a natural product used to treat leishmaniasis (11, 12). It is a polyene antimicrobial and was initially introduced for antifungal use alongside other polyenes, such as nystatin (Nys) (13–21). The introduction of liposomal formulations of AmB, AmBisome^®^, significantly reduced inherent toxicity (12,22–24) and has become the preferred treatment for VL (8,25,26), especially while costs have been kept low due to donations of the drug to the WHO for use in countries most affected by the disease (27–32). AmB is used in the elimination program of VL in south Asia (1) and has remained effective in VL patients refractory to other antileishmanials (33). It is also active against other forms of the disease (34, 35), although it has been used less for CL (12, 36). The availability of donated AmBisome^®^ has enabled a campaign of single dose use of the drug. Treating large numbers of individuals in this way avoids complications associated with multiple-dosing regimens and has been of public health benefit (37–39), contributing to a decline in infections as part of a WHO led campaign to eliminate VL as a public health problem (40, 41). That campaign, however, has assumed that the development of resistance to AmB in *Leishmania* parasites is not a significant problem. This is based on the widely held perception that, despite over 70 years of clinical use as an anti-fungal agent, this use has not been accompanied by a significant emergence of AmB resistance (AmBR) in fungal species (42–49). AmB, and other antimicrobial polyenes, bind to ergostane-based membrane sterols with high avidity, and ergosterol and related sterols are the predominant forms in fungi and also in *Leishmania* spp. (50–55). The compounds bind to cholesterol, the major sterol in mammalian cell membranes, with markedly less avidity (56). This selectivity is driven by the altered structural configuration of ergosterol and its analogues relative to cholesterol: differences include double bonds at positions C7 and C22, and a methylation at C24 of the side chain (57–59).

In spite of the perception that AmBR is only a minor risk, several examples of resistance (60–65) and treatment failure with AmB (47,66–68) have been described in leishmaniasis cases, and polyene resistance is by no means unknown in fungal disease (49,64,69–79). An outbreak of autochthonous canine VL in Uruguay found clinical isolates 3 to 4-fold less susceptible to AmB (80, 81). Similar differences in susceptibility to AmB were found in a non-endemic region in India (63) and also patients with cutaneous leishmaniasis in Colombia (82).

Development of AmBR is a major threat for the treatment of VL (83). Mechanisms of resistance appear to be diverse. For example, fungal species with a normal content of ergosterol and high levels of catalase are intrinsically resistant to AmB (45,84,85). Furthermore, in *Leishmania* spp., numerous enzymes involved in the protecting against reactive oxygen species (ROS) appear to associate with AmBR. These include the polyamine-trypanothione pathway (PTP) (60,86,87), L-asparaginase (88), cysteine synthase (89), ascorbate peroxidase (90), tryparedoxin peroxidase (91) and the silent information regulator 2 (Sir2) protein (92). Furthermore, cross resistance to AmB is sometimes observed between lines resistant to the antileishmanials miltefosine and antimonials (13,65,93,94), reflecting a multifactorial mode of action (MoA) of AmB (43, 60).

We have previously shown that ergosterol loss relates to selection of AmBR in laboratory-generated mutants of *Leishmania mexicana*. Genome sequence analysis of resistant mutants has shown that mutations in genes encoding lanosterol 14-alpha demethylase (C14DM) (95), C5-sterol desaturase (C5DS) and C-24 sterol methyl-transferase (C24SMT) (96) can all lead to AmBR. Additionally, sterol profiling revealed loss of ergosterol with inferred loss of drug binding in each case. Other studies in AmBR *Leishmania* spp. have also shown loss of expression of one of the two C24SMT-transcripts (60,97–99) and structural variations at this locus, alongside loss of function of the miltefosine transporter in some instances (93,100– 102). Moreover, using *L. major* and *L. donovani*, a series of gene knockout studies have also shown that loss of C14DM (103–105), C24SMT (99, 106) and C5DS (107) all lead to the creation of viable and AmBR-parasites, albeit of varying degrees of fitness. Here we describe the characterisation of 15 new independently selected polyene-resistant lines of *L. mexicana* (ten selected to AmB and four to Nys), plus one AmBR-*L. infantum* line. In each case sterol profiling pointed to a loss of ergosterol and its analogue with essentially two different types of compensatory-profile arising, which are each accounted for by mutations to mutations in C24SMT or C5DS. We also assessed the fitness of the selected lines in mice and stability of the resistance phenotype.

## Results

### Polyene resistance selection in fifteen independent *Leishmania* spp. lines

A total of fifteen independent lines of *Leishmania* spp. (S1 Table) were selected for resistance via a stepwise increasing concentration (S1 Fig) of the polyenes AmB or Nys being added to the culture media HOMEM or a serum-free defined medium (DM hereon) (S2 Table). Seven AmB resistant *L. mexicana* lines (AmBRcl.14, AmBRcl.8, AmBRcl.6, AmBRcl.3, AmBRA4, AmBRB2 and AmBRC1) were selected in complete HOMEM and three AmB resistant lines (AmBRA3-DM, AmBRB2-DM and AmBRC2-DM) were selected in DM. In addition, four Nys resistant (NysR) *L. mexicana* lines (NysRcl.B2 NysRcl.C1, NysRcl.E1, and NysRcl.F2) and one AmBR *L. infantum* (AmBRcl.G5) were also selected in complete HOMEM. The levels of resistance attained in *L. mexicana* after AmB exposure for 210 days in HOMEM were between 4.5- and 12.8-fold compared with the wild type cultured in parallel in the absence of drug. A comparable increase in resistance was observed in all four lines exposed to Nys for 140 days ranging from 6.6- to 11-fold and to a lesser extent in AmBRcl.G5 (*L. infantum*) which increased by 3.7-fold after drug pressure for 105 days. In contrast, resistance attained in all three AmBR *L. mexicana* lines selected in DM over 372 days was only 2.2-fold that of the wild type in DM. Levels of resistance to AmB or Nys were stable after at least 10 passages in the absence of drug and confirmed in at least two clonal populations derived by limiting dilution from each of the independent fourteen polyene resistant *L. mexicana* lines (for AmBR-*L. infantum*, only one clonal population could be recovered and analysed). AmB can be sequestered by cholesterol and other lipoproteins present in the serum (108, 109). Surprisingly, wild type cells sustained in serum-free conditions (DM) were 3.6-fold less susceptible to AmB (EC_50_ 217.3) than they were in HOMEM (EC_50_ 60 nM) indicating an enhanced overall potency of the drug against parasites in rich medium. Parasite growth rate of AmBR and NysR resistant lines cultured in HOMEM was similar to the parental line cultured in parallel without drug whilst in DM AmBR resistant lines was slightly lower (S2A-C Fig). Interestingly, four AmBR lines sustained in HOMEM alongside the parental line showed no growth after switching from HOMEM to DM (S2D Fig). All the resistant cells were morphologically comparable to wild type (WT) with the exception of AmBRcl.14 and AmBRcl.3 that had an increased cell body length relative to wild type (S2E Fig).

### Response of polyene resistant lines to other antileishmanials

Susceptibility to other antileishmanials was measured in only one resistant clonal population derived from each of the fifteen lines by limiting dilution (S1 Table). Considering the similar mode of action between polyenes, we expected reciprocal cross-resistance between AmBR lines to Nys and *vice versa*. AmBR lines were 12.9 to 25-fold cross resistant to Nys. To a lesser extent (1.8 to 2.0-fold), cross-resistance was also apparent with the smaller polyene natamycin (NMC). Similarly, NysR lines cross-resistance to AmB was from 5.8 to 9.1-fold. Variable cross resistance to miltefosine (MF) was found across the polyene resistant lines with the lowest change found in lines cultured in DM (1.1 to 1.4-fold) while cells selected in HOMEM showed changes ranging from 2.3 to 5.3-fold increases in most lines, and, rising to 10-fold in two lines with a deletion of the miltefosine transporter (described in a later section) AmBRcl.8 and AmBRcl.6 (Fig 1). In previous studies, this locus has been associated with cross resistance between AmB and MF (93,100–102). A range of changes in EC_50_ were obtained with two inhibitors of lanosterol 14-alpha demethylase (C14DM), ketoconazole and fenarimol (103,110–113), and to potassium antimony tartrate (PAT). All fourteen polyene-resistant *L. mexicana* lines cultured in HOMEM and DM were significantly more susceptible to paromomycin (PAR) (P=0.0001 to 0.0319), with a decrease of 1.7 to 10-fold in the EC_50_ irrespective of the culture medium (Fig 1, S3 Table and S3 Fig). Similarly, significant hyper-susceptibility to pentamidine (PENT) and to methylene blue (MB) was found in all polyene resistant lines.

**Fig 1.**
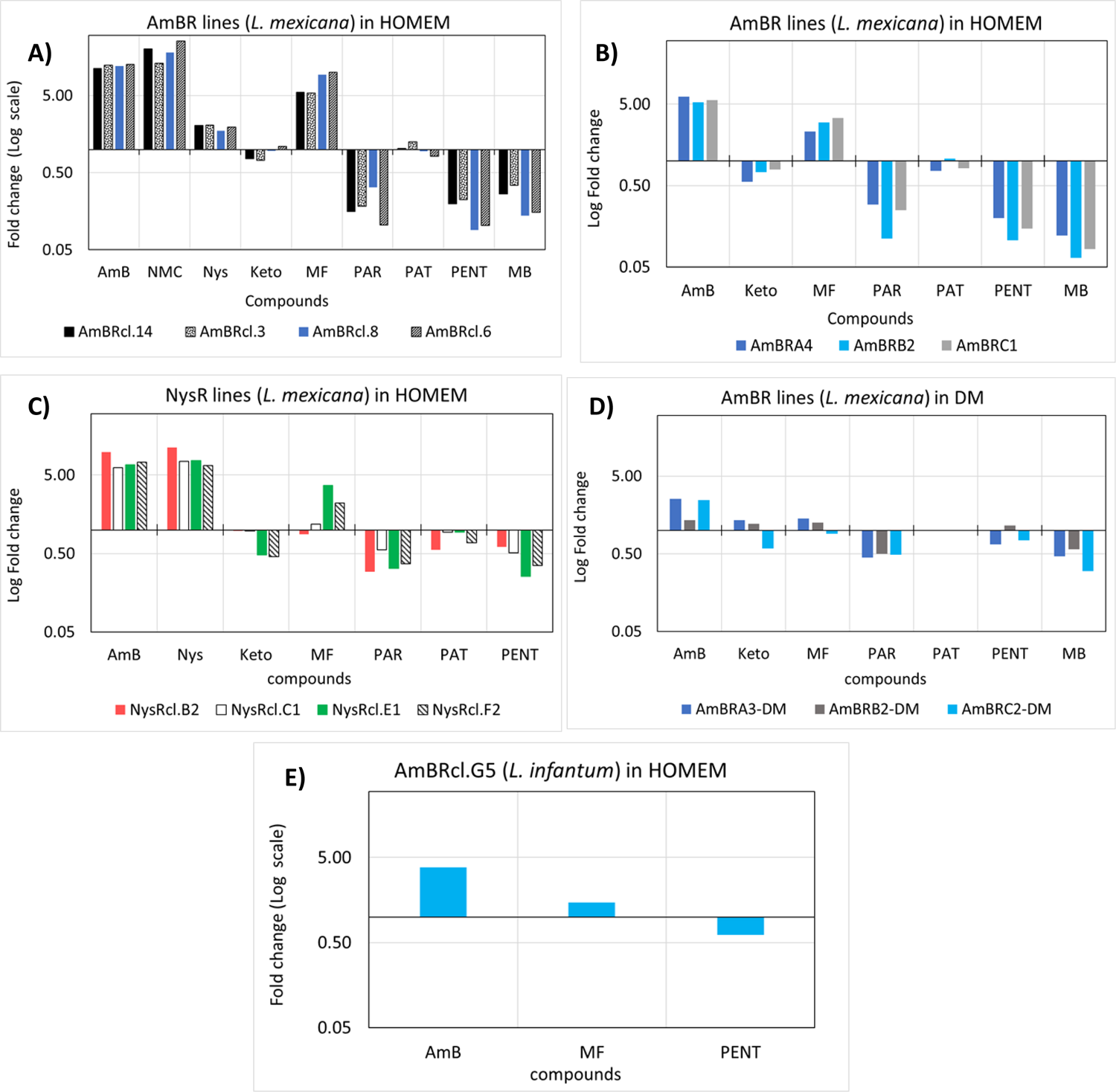
Fold change in EC_50_ in all polyene resistant lines of *Leishmania spp*. Log Fold change (EC_50_) in resistant lines is relative to the respective parental wild type cultured in parallel without drug. **A-B)** AmBR-*L. mexicana* in HOMEM, **C)** NysR-*L. mexicana* in HOMEM, **D)** AmBR-*L. mexicana* in DM and **E)** AmBR-*L. infantum* in HOMEM. Values higher and lower than 1, indicate a decrease and rise in susceptibility relative to the parental line, respectively. A complete list of values in fold-change and EC_50_ with statistical analysis is provided in S3 Table and S3 Fig. AmB (amphotericin B), Nys (nystatin), Keto (ketoconazole), MF (miltefosine), PAR (paromomycin), PAT (antimony potassium tartrate), PENT (pentamidine) and MB (methylene blue).

EC_50_ value to PENT decreased by 7- to 10.9–fold (P=0.0008 to P<0.0001) in all seven AmBR lines and from 2- to 3.8-fold (P=0.01 to P<0.0001) in all four NysR lines, selected in HOMEM (Fig 1, S3 Table and S3 Fig). Recently, Mukherjee et al. related changes in sterol composition to altered mitochondrial membrane potential (ΔψM) which they implicated in hypersensitivity to PENT whose uptake into mitochondria depends on ΔψM (105). To a lesser extent, AmBR lines grown in DM also revealed hyper-susceptibility to PENT (P=0.0938 and P=0.2143), except AmBRB2 which was marginally less sensitive (1.15-fold) to the drug. It is possible the differences in oxidative stress induction by AmB in different media also relate to this difference in PENT sensitivity between resistant lines selected in different media. A comparable rise in susceptibility to PENT (P ≤ 0.05) was also observed in clemastine fumarate-resistant *L. major* promastigotes with altered membrane lipid composition (114), further indicating potential trade-offs between AmB and PENT resistance mechanisms, centred around lipid production. Exposure to MB, an oxidative stress inducing agent (115), showed that AmBR lines grown in HOMEM were 2.9 to 15.3-fold more susceptible than wild type cells (P= 0.0001 to 0.0084) to this agent which induces intracellular oxidative stress by increasing the NADP+: NADPH ratio and stimulating the oxidation of key cellular redox related thiols such as glutathione or its analogue trypanothione in trypanosomatids (116, 117). To a lesser extent (1.76 to 3.34-fold), hyper-susceptibility to MB was also found in AmBR lines selected in DM (AmBRA4, P=0.506; AmBRB2, P=0.114 and AmBRC1, P=0.012) (Fig 1D, S3 Table and S3 Fig). Increased susceptibility to oxidative stress was observed in other AmBR lines exposed to glucose oxidase which creates hydrogen peroxide in medium (95, 101).

### Metabolic response to AmB exposure

AmB kills *Leishmania* very quickly. At high concentration of AmB (i.e., 5 x the EC_50_) death ensues within 30 min. We used untargeted metabolomics to determine what changes occur to cellular metabolism over 15 mins by exposing parasites to high concentrations of AmB equivalent to 5x their respective EC_50_ values in HOMEM AmB: 0.31 μM (parents LmWT1 and LmWT2); 3.0 μM (AmBRcl.14 and AmBRcl.8) and 1.74 μM (AmBRB2). Likewise, cell lines cultured in DM were subjected to AmB at high concentrations (i.e., 5 x EC_50_): 1.16 μM (LmWT-DM) and 3.58 μM (AmBRA3-DM). In untreated cells no significant differences were seen in intracellular metabolites involved in the tricarboxylic acid cycle (TCA) or the PPP between HOMEM and DM (S4 Table).

We then compared the metabolomes of cells exposed to drug that had been cultured under the differing media conditions. The most notable difference was in the major increase in metabolites of the pentose phosphate pathway (PPP): metabolites such as 6-phospho-D-gluconate (Adj. P-value = 0.0121 in AmBRcl.14; 0.0063 in AmBRcl.8; 0.0221 in AmBRB2 and 0.0074 in AmBRA3-DM), glyceraldehyde-3-phosphate (Adj. P-value = 0.1377 in AmBRcl.14; ND in AmBRcl.8; 0.0182 in AmBRB2 and 0.0012 in AmBRA3-DM), sedoheptulose7-phosphate, (Adj. P-value = 0.6936 in AmBRcl.14; 0.0026 in AmBRcl.8; 0.0099 in AmBRB2 and 0.0003 in AmBRA3-DM), ribose 5-phosphate (Adj. P-value = <0.0001 in AmBRcl.14; ND in AmBRcl.8; 0.0016 in AmBRB2 and ND in AmBRA3-DM) and D-glucono-1,5-lactone 6-phosphate (Adj. P-value = 0.0002 in AmBRcl.8; 0.0057 in AmBRB2, not detected in other lines) were significantly increased in abundance in cells exposed to drug in HOMEM (Fig 2A) but not in DM medium (Fig 2B). Upregulation of the PPP (Fig 2C) points to an increased oxidative stress response (119–121). This difference may explain why HOMEM-cultured cells are more sensitive to drug and to the oxidative stress inducer methylene blue (Fig 1 and S3 Table), with these cells experiencing heightened oxidative stress over and above that induced through cellular lysis following pore formation (triggered via AmB-ergosterol binding). This observation may also explain why the fold-change in resistance of the lines selected in HOMEM (5.29 to 12.79-fold) is greater than those achieved in DM (1.36 to 2.54-fold) (Fig 1 and S3 Fig), as alterations in oxidative stress response may greatly enhance HOMEM-based resistance.

**Fig 2.**
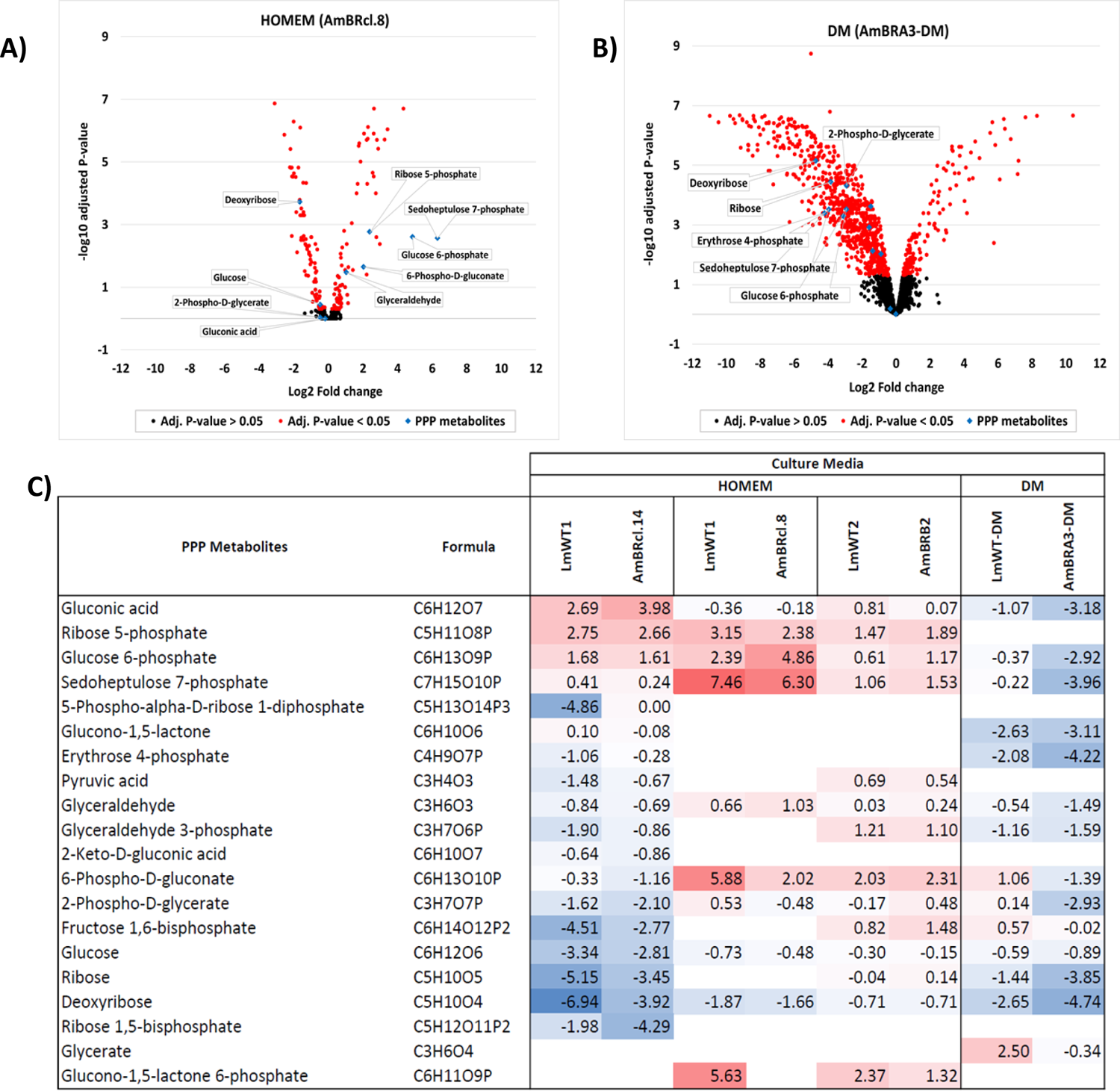
Metabolic effects of exposure to AmB in *L. mexicana* promastigotes cultured in HOMEM and DM. Mid log AmBR lines alongside with their parental wild-type *L. mexicana* promastigotes (1 x 10^8^) were cultured in HOMEM (LmWT1, AmBRcl.14, AmBRcl.8, LmWT2 and AmBR2) and DM (LmWT-DM and AmBRA3-DM). **A)** Volcano plot showing significant log2 fold-changes (red dots) of peaks as detected by LC-MS in promastigotes cultured in HOMEM (AmBRcl.8) and **B)** in DM (AmBRA3-DM). Multivariate data analysis was performed with PiMP analysis pipeline (118) and the Benjamini-Hochberg procedure adjusted raw P-values (q-values) < 0.05 for ANOVA. **C)** Heat map of representative intermediates of the PPP. Each line represents one biological replicate (n=4). Increase (red) or decrease (blue) in fold-change is relative to untreated cells after treatment with AmB at 5 x their EC_50_ values for 15 mins. Values in bold indicate metabolites identified and all others are putative based on mass. Empty boxes (no value) indicate that the metabolite was not detected. A full list of P-values of the metabolome (including PPP intermediates) is provided in S4 Table.

### Sterol profiling in polyene resistant lines of *Leishmania* spp

Binding to ergosterol is the primary event in the mode of action of antimicrobial polyenes (43, 122). Previous studies with AmBR *Leishmania* spp. have shown that loss of ergosterol commonly accompanies resistance development (60,97,101,123–126). Fig 3 shows the sterol biosynthesis pathway (SBP) in *Leishmania* adapted from the KEGG database (https://www.genome.jp/kegg/) and previous work (101,127–129).

**Fig 3.**
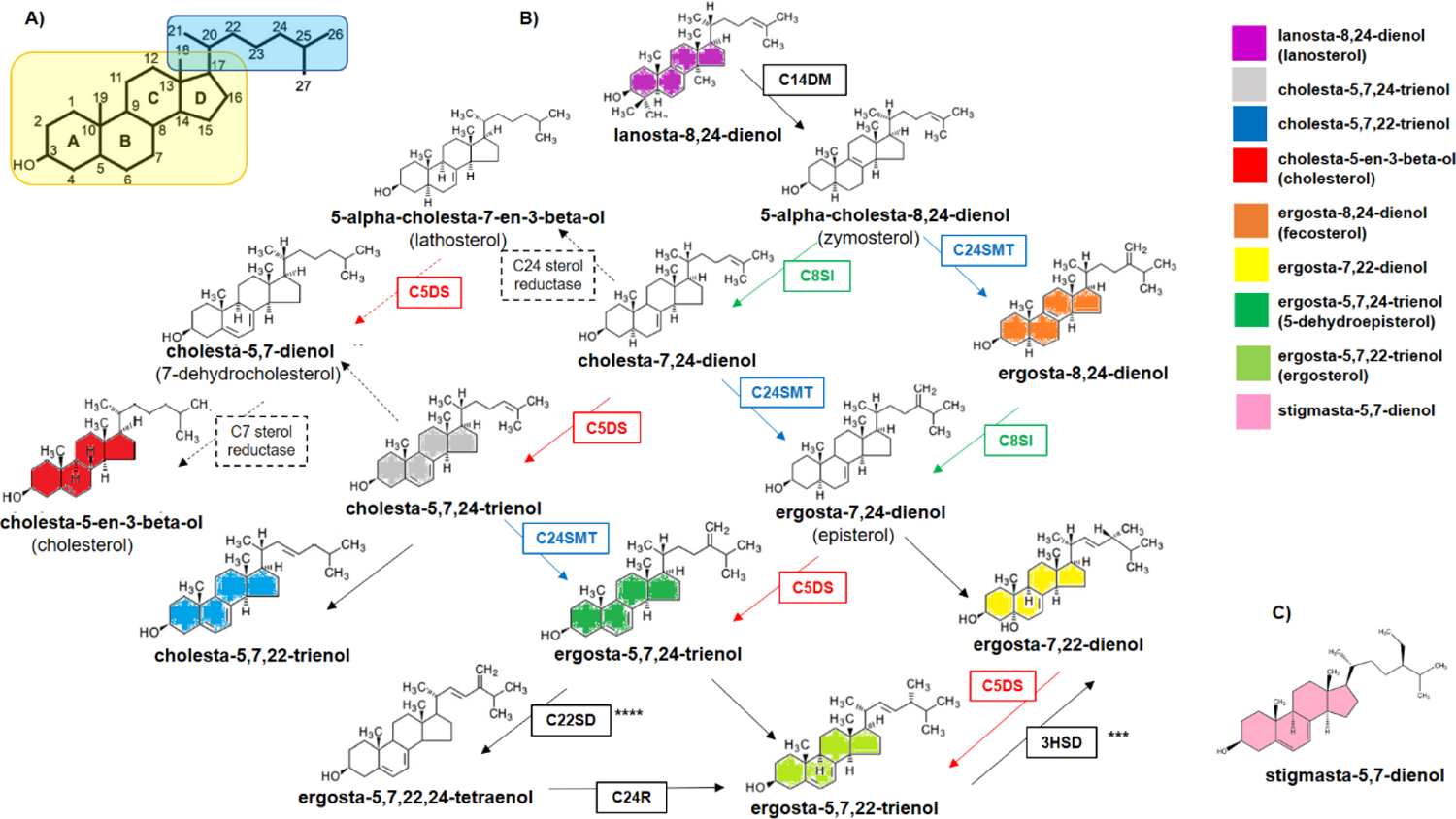
Sterol Biosynthesis in *Leishmania spp*. **A)** Numbering system of the carbons in the sterol ring (boxed in yellow) and side chain (boxed in blue). **B)** Molecules coloured indicate sterols found in this work. Enzymes identified in *Leishmania spp*. (solid arrows / boxes) are sterol C14 demethylase*, LmxM.11.1100* (C14DM); sterol C24-methyltransferase, *LmxM.36.2380* and *LmxM.36.2390* (C24SMT) (blue); C8-sterol isomerase, *LmxM.08_29.2140* (C8SI) (green); sterol C5 desaturase or lathosterol oxidase, *LmxM.23.1300* (C5DS) (red); sterol C22 desaturase, *LmxM.29.3550* and/or *LmxM.33.3330* (C22SD); sterol C24 reductase, *LmxM.32.0680* (C24R) and 3-beta hydroxysteroid dehydrogenase, *LmxM.18.0080* (3HSD). Also shown some enzymatic steps known in mammals (dotted arrows / boxes). **C)** The position of stigmasta-5,7-dienol within the pathway is unknown. Modified from: KEGG Database (https://www.genome.jp/kegg/); Pountain et al, 2019; Yao & Wilson, 2016 (***) and Bansal et al., 2019 (****).

The identities and composition of the sterol peaks detected by gas chromatography-mass spectrometry (GC-MS) based on matches with the standards used and with the NIST library are shown in S4 Fig & S5-A Table (https://www.nist.gov/srd/nist-standard-reference-database-1a-v14). Ergosta-5,7,24-trienol (C_28_H_44_O) was the most abundant sterol detected here in wild type *Leishmania* spp. promastigotes ranging from 72 to 82% of total sterol (Fig 4). Also known as 5-dehydroepisterol, this isomer of ergosterol differs from the major fungal ergosterol, ergosta-5,7,22-trienol (128) in the position of the double bond (carbon 24 rather than carbon 22) and in its chromatographic retention time (S5-A Table) (103, 130).

**Fig 4.**
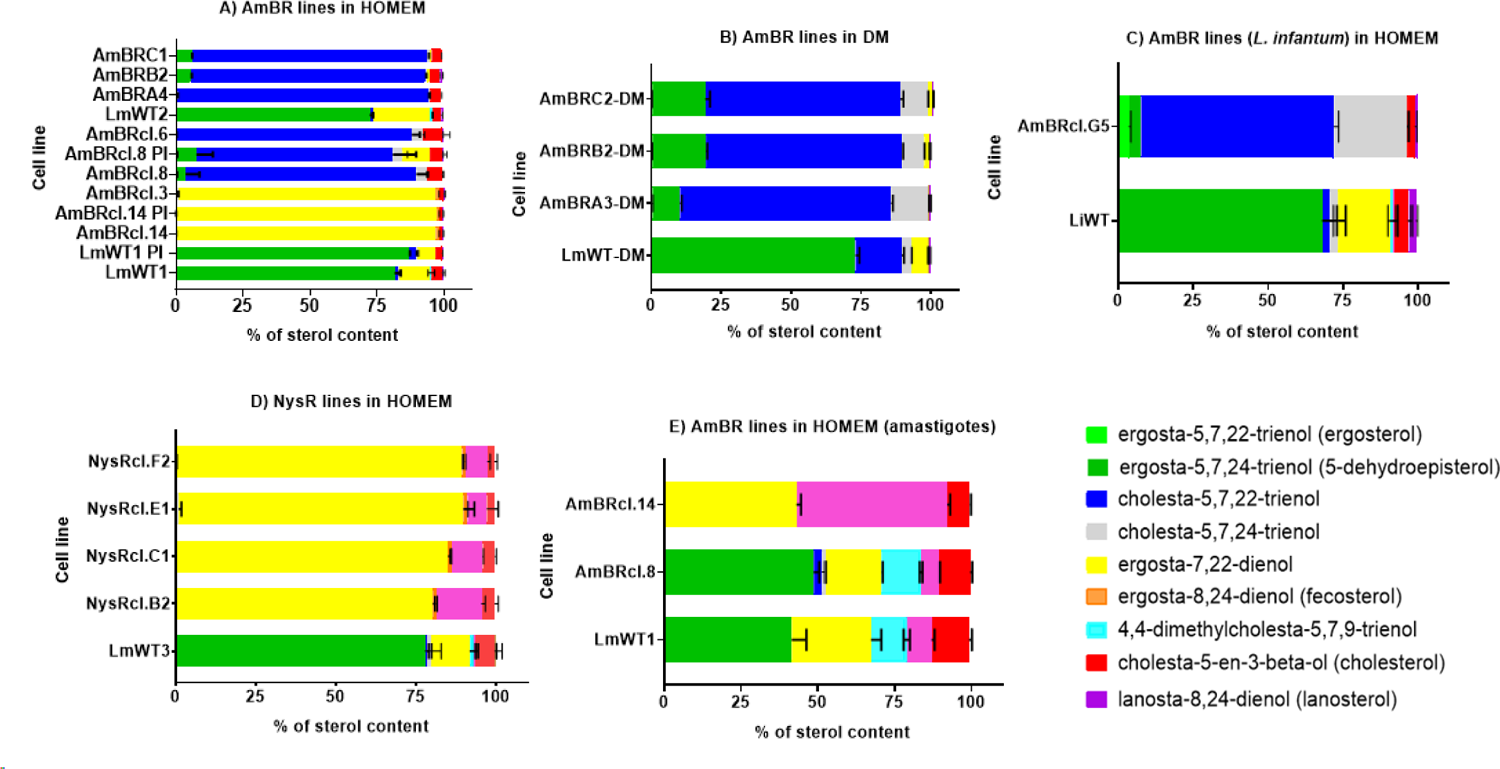
Percentage of sterols (GC-MS) in polyene resistant lines of *Leishmania spp*. Standard deviation of the mean (bars) is from three biological replicates. Parasites were cultured as promastigotes in HOMEM (panels A, C and D) and DM (panel B) or as axenic amastigotes (recovered after infection in mice) in Schneider’s media (panel E). Species are *L. mexicana* (panels A, B, D and E) and *L. infantum* (panel C) and lines are resistant to polyenes AmB (panels A, B, C and E) and Nys (panel D). See S5 Table and Material and Methods for a full description of the content and identification of sterols with GC-MS.

Another abundant sterol in wild type promastigotes was ergosta-7,22-dienol (11%) followed by cholesterol (2 to 7%) (Fig 4). Cholesterol was absent in all AmBR and wild type parasites cultured in DM (Fig 4D), which is unsurprising given that cholesterol is derived from exogenous sources, including the serum added to culture media (131, 132) and from the membrane of the host-macrophage (133). This possibly explains its higher abundance in the intracellular stage of the parasite in both wild type (12.2%) and AmBR lines (7.3 to 10.2%) cultured in HOMEM (Fig 4E & S5-B Table). All of the polyene-selected lines had altered sterol profiles compared to wild-type and these fell into two broad types. In three of the AmBR *L. mexicana* lines selected in HOMEM and all four of the Nys resistant *L. mexicana* lines, the predominant sterol became ergosta-7,22-dienol (96 to 97%) (Fig 4A & D), which lacks the C5 desaturation of AmB-binding ergosterol. Another sterol, stigmasta-5,7-dienol, was also present in all NysR lines, more abundant in NysRcl.B2 (9.9%), NysRcl.C1 (14.2%), NysRcl.E1 (6.1%) and NysRcl.F2 (7.0%).

We also analysed the sterol content of axenic amastigotes in two resistant lines alongside with their parental wild type following recovery from mouse tissue (post-infection) parasites were maintained as promastigotes in HOMEM and transformed into amastigotes in Schneider’s medium. Interestingly, stigmasta-5,7-dienol was also present in wild type *L. mexicana* axenic amastigotes (8.1% of total) and unchanged or little depleted in AmBRcl.8 (5.8%) while increased notably in AmBRcl.14 (49.1%) (Fig 4E). However, its role in resistance and its position within the sterol biosynthetic pathway awaits further investigation (Fig 3C). In five of the HOMEM selected *L. mexicana* AmB lines and all of the AmB lines selected in DM, as well as the *L. infantum*, cholesta-5,7,22 trienol became predominant. In cells selected in HOMEM it comprised 86%-93% of the total sterol, while in those cells selected in DM this proportion was somewhat lower (69.7%-75.4%) whilst an isomer, cholesta-5,7,24-trienol (C27H42O) increased from 3.4% abundance in wild-type to 7.9%-12.3% in resistant lines selected in DM. In AmBR *L. infantum* 64% of the total comprised this sterol while a further 25% of the total was its isomer cholesta-5,7,24 trienol. Interestingly, cholesta-5,7,22-trienol was higher in wild type cells sustained in DM (16.5%) in comparison with values found in all parental lines cultured in HOMEM (0.5 to 2.57%) (Fig 4A-C and S5-B Table).

### Genomic changes identified with whole genome sequencing in polyene-resistant Leishmania spp. lines

Whole genome sequencing was performed on each of the 15 polyene resistant cell lines and particular attention paid to gene deletion or amplification and SNPs found repeatedly in the same gene in different isolates. Given the altered sterol profiles reported in the previous section, we also systematically inspected all genes encoding enzymes of the sterol pathway. All polyene resistant *Leishmania* lines analysed here revealed mutations to one or other of two genes, sterol C5-desaturase (C5DS) and sterol C24-methyl transferase (C24SMT), encoding enzymes of the sterol pathway.

### Sterol C5 desaturase mutations

One group of three resistant lines revealed point mutations to C5DS (or lathosterol oxidase (LOX), gene ID = *LmxM.23.1300*). In AmBcl.3, all of the reads of C5DS are mutated (S4 Fig) containing one of two independent heterozygous mutations within the same codon, G220A (V74M) and T221A (V74E), leading to a substitution of a non-polar valine residue, which is conserved across the aligned *Leishmania* species (S5 Fig), by a polar methionine and a negatively charged glutamate residue, respectively. A second homozygous substitution G731T (R244L) was present in AmBRcl.3 and AmBRcl.14 (Fig 5A). Interestingly, both lines also showed a homozygous variant G208A (G70S) that changed a non-polar glycine into a polar serine residue in the adjacent gene downstream from C5DS, a dynein heavy chain (ID = *LmxM.23.1310*) (Fig 5B). Also, in C5DS, AmBRcl.14 showed an in-frame deletion ATG277-279 (M93del) while in NysRcl.B2 a neighbouring frame-shift deletion G286del (A96del) was present (Fig 5A-B), leading to loss of a full-length coding region in each case. In all these C5DS mutants an accumulation of ergosta-7,22-dienol (Fig 4) was observed, indicating that all of these different mutations impacted upon activity of the enzyme leading to an accumulation of its substrate whose use is ablated in the mutants.

**Fig 5.**
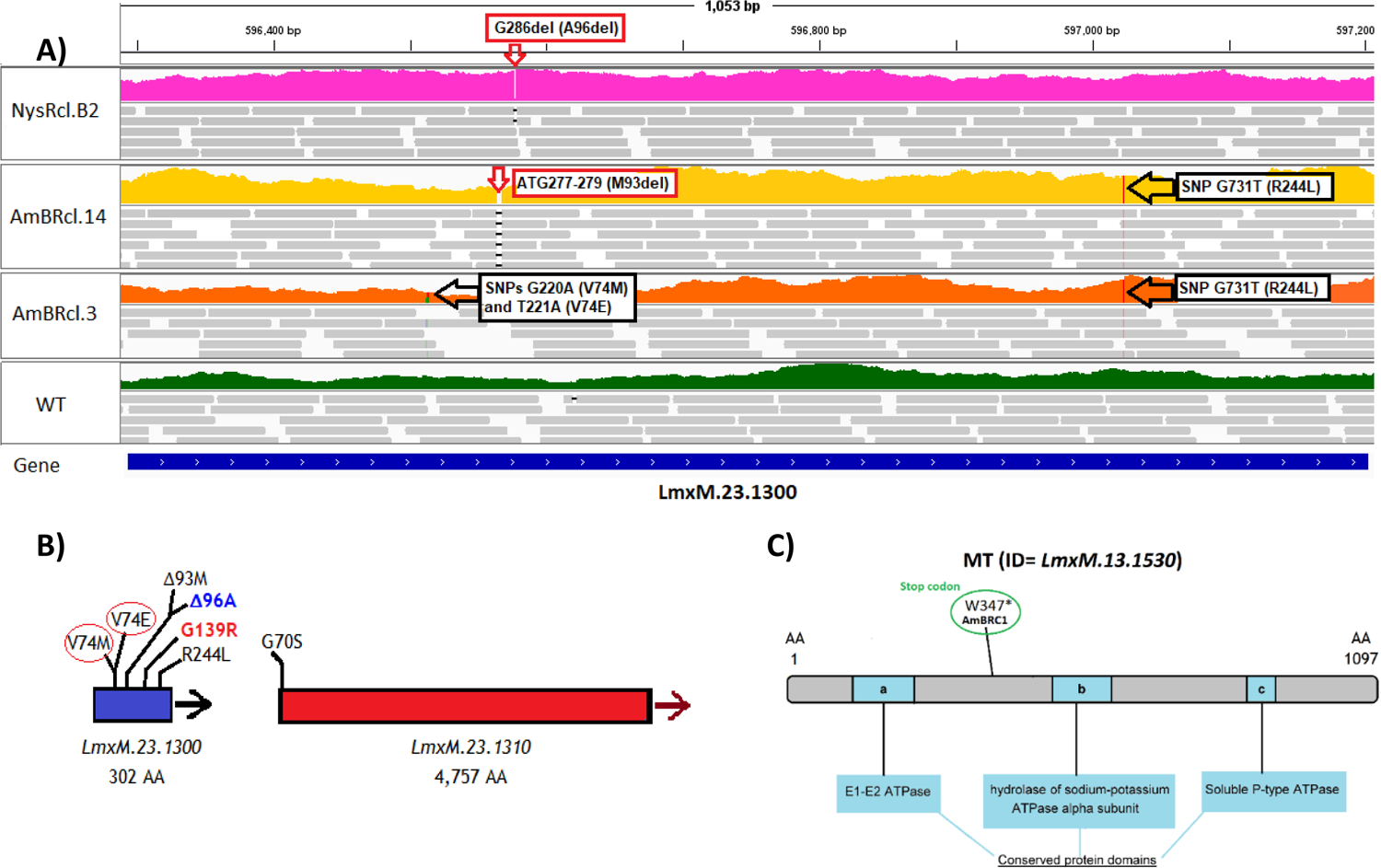
Mutations in lipid-associated genes found in polyene resistant lines of *L. mexicana*. **A)** Coverage is shown in colours for NysR (top in pink), AmBRcl.14 (upper-middle in yellow), AmBRcl.3 (lower middle in orange) and wild type (WT) (bottom in green) and reads (grey bars) with mapping quality, > 0 are shown for each line. The blue line at the bottom indicates the open reading frame of the C5DS (or LOX) *LmxM.23.1300* gene and variants are indicated (empty arrows). Image produced with WGS data aligned to the reference genome (https://tritrypdb.org) using IGV_2.8.9 (http://software.broadinstitute.org); **B)** Cartoon showing the variants linked to polyene resistance in LOX (C5DS) (blue bar), a SNP in the gene downstream (*LmxM.23.1310*) is also shown (red bar). The red circles indicate that these two mutations are in the same codon and are independent (found in different reads). Mutations found in AmBRcl.14 and AmBRcl.3 (in black) and in Nyscl.B2 (blue) are shown alongside with a variant G139R (red) reported in our study (Pountain et al, 2019) which indicates a residue involved in catalysis or ligand binding; **C)** Cartoon of MT gene *LmxM.13.1530* portraying conserved domains and the proximity of the variant W347* (stop codon) found in this study in AmBRC1 (green circle).

### Sterol C24 methyl transferase mutants

The second sterol pathway gene that was mutated in multiple separate lines was the C24SMT tandem array (*LmxM.36.2380* and *LmxM.36.2390* in *L. mexicana*) (S6 Fig & Table 1). Five AmBR lines of *L. mexicana* cultured in HOMEM, AmBRcl.8, AmBRcl.6, AmBRA4, AmBRB2 and AmBRC1 showed no coverage of the intergenic region (∼2.5 kb) between the two copies (S6A-B Fig). The *L. infantum* AmB resistant line (AmBRcl.G5) also showed a loss of coverage across this intergenic region (S6E Fig). Copy number variation (CNV) confirmed a deletion within the C24SMT tandem pair in all of these lines (S8 Table). In AmBRA4, a novel non-silent mutation A325V (C974T) resulted in a substitution from alanine to valine. In all of the AmB resistant lines selected in DM, this intergenic region was not of decreased coverage (S6C Fig), but a variant G961A (V321I) which constitutes a locus that differs between the two copies of C24SMT (Pountain et al, 2019) was found. All of the lines where mutations were targeted around the C24SMT locus demonstrated an increase in cholesta-5,7,22-trienol (Fig 4A-C), which is the substrate for C24SMT, indicating that in each case the mutations have diminished the activity of that enzyme. This locus showed no genomic changes in NysRcl.B2 (S6D Fig), in which the predominant sterol was ergosta-7,22-dienol (Fig 4 and S5-B Table) compatible with a mutation in C5DS as noted above.

**Table 1.**
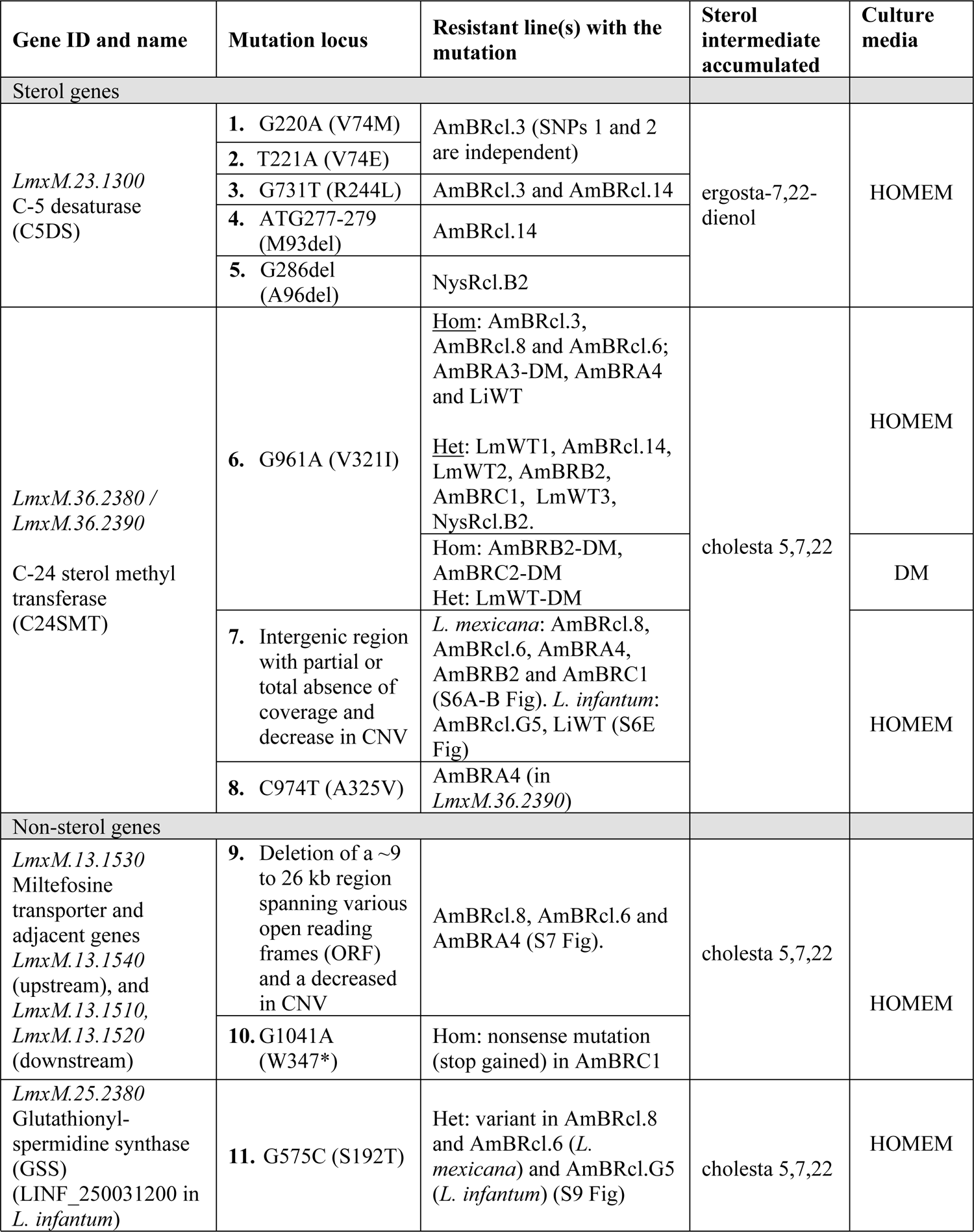
WGS in sterol and non-sterol genes in polyene resistant lines of *Leishmania* spp. Gene IDs were retrieved from TriTrypDB in *L. mexicana*. Genotype and amino acid changes from variants in sterol genes is provided in S6 Table. A complete list of variants is provided for SNPs (S9 Table) and CNVs (S8 Table). Hom: homozygous; Het: Heterozygous.

### Mutations to the Miltefosine transporter and genes involved in oxidative stress defence

Several previous studies on resistance to AmB (93, 101) showed that the miltefosine transporter gene (*LmxM.13.1530*) was also deleted during the process of selection of AmB resistance. Here we noted that in three lines (AmBRcl.8, AmBRcl.6 and AmBRA4) the MT and an adjacent gene (*LmxM.13.1540*) were deleted (S7C-D Fig). In AmBRA4 the deleted region was larger (∼20 kb) spanning two additional genes (S7D Fig). CNV analysis confirmed that the MT gene was, however, retained in all of the other lines. A reduction in CNV in this locus was also found in AmBRcl.14, AmBRcl.3 and NysRcl.B2 (they lost one copy of the two) but increased in AmBRB2. Changes in ploidy in all lines are shown in S8 Fig and S8 Table). In all of the lines selected in DM, the MT gene remains (S7E Fig), but in one line, AmBRC1, a homozygous variant G1041A (W347*) appears, indicating a translation termination (stop codon) and consequent loss of function of MT that resulted in 3.3-fold resistance to MF relative to WT (Fig 5C & Table 1).

Other genes beyond those encoding members of the sterol pathway were also found in two or more polyene-resistant lines and these are listed in Table 1, S10 Fig and S9 Table and their roles or actual relationship with resistance awaits further analysis, although it is notable that four genes encoding enzymes associated with the leishmania thiol redox system (S9B Fig) were found. Notably, these mutations are present in cells selected in HOMEM medium in which oxidative stress was more profound. In AmBRcl.G5 (*L. infantum*), a heterozygous non-synonymous mutation G575C (S192T) (S9A Fig) was present in the glutathionylspermidine synthase (GSS) encoding gene (ID = *LINF_250031200*, formerly annotated as *LinJ25.2500*). Strikingly, the same residue G575C (S192T) in GSS (*LmxM.25.2380*) was also mutated in two *L. mexicana* lines, AmBRcl.8 and AmBRcl.6 (S9C Fig) both which showed an attenuated phenotype in mice (Fig 6-A). Moreover, in five lines, CNV revealed relatively small changes in another four genes involved in oxidative stress, two copies of the type II tryparedoxin peroxidase (glutathione peroxidase-like) (*LmxM.26.0800* and *LmxM.26.0810*), glutaredoxin (*LmxM.26.1540*) and glutathione S-transferase (*LmxM.12.0360*). Previous work has pointed to a potential role of the leishmanial redox system (S9B Fig) in resistance to AmB (60,87–89,91,92,134). The actual role of these genes awaits further investigation, but it seems likely that changes to the redox system might indeed be important particularly during selection in HOMEM where higher levels of oxidative stress in cells exposed to AmB are manifest.

**Fig 6.**
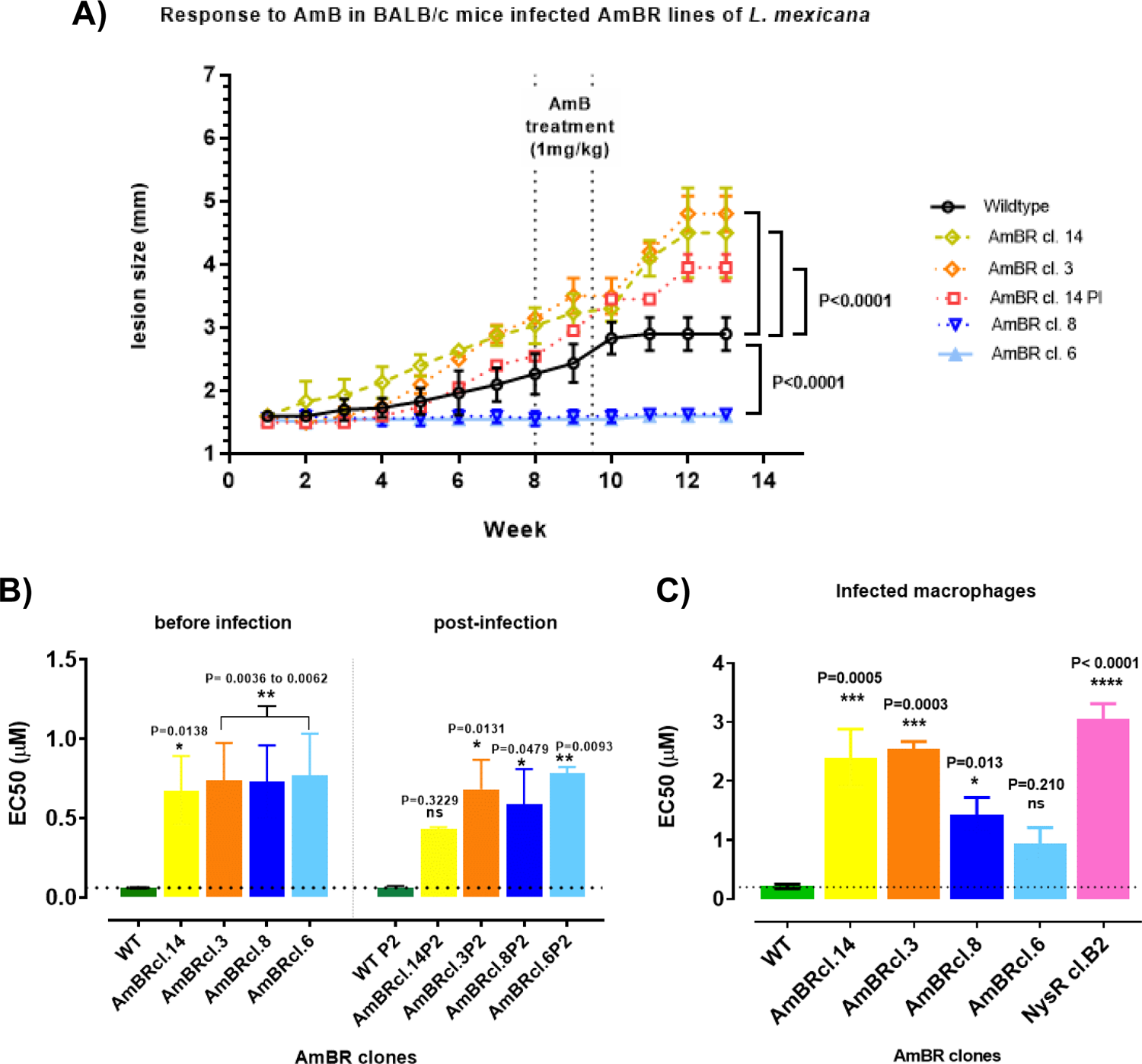
AmB activity in polyene resistant *L. mexicana*. **A)** Stationary promastigotes were inoculated at 2 x 106 into 500 µl of PBS. Evolution of lesion was measured weekly. BALB/c female mice were two-months old at the time of inoculation. Treatment: 1 mg per kg IV every other day in the tail vein with a total of six injections (∼120 µl) of AmB (deoxycholate). **B)** activity of AmB in axenic promastigotes before and after infection of mice; **C)** AmB activity against AmBR- and NysR-*L. mexicana* in infected macrophages. Values (µM) are the mean EC50 ± SD (bars) (See S7 Table). Tukey’s multiple comparison test was used to find pairwise differences between resistant lines and parental wild type. Statistically significant values (P<0.05, 95% confidence interval) are indicated with stars as follows: P>0.05; * P≤0.05; ** P≤0.01; *** P≤0.001; **** P≤0.0001).

### Infectivity of AmBR lines *in vivo*

We initially selected AmB resistance in cultured promastigotes given the tractability of this system for biochemical analysis. However, it is important to understand whether findings from promastigotes translate into amastigote forms that replicate in mammalian macrophages and which are associated with disease. Therefore, we chose four of the AmBR mutants to test their ability to retain infectivity and virulence in mice. Two of these were from the C5DS mutant group (AmBRcl.14 and AmBRcl.3) and two from the C24SMT mutant group (AmBRcl.8 and AmBRcl.6), in these groups, the wild type sterol (ergosterol) was replaced by ergosta-7,22-dienol and cholesta-5,7,22-trienol, respectively. For these experiments, we also included a low passage (P2) isolate (AmBRcl.14 PI) recovered from a previous infection with AmBRcl.14. Notably, mice infected with AmBRcl.14 (C5DS), AmBRcl.3 (C5DS) and AmBRcl.14 PI (C5DS) showed an increase in the size of the lesion at the inoculation site (footpad) up to two times greater (week thirteen) than that observed in mice infected with the parental wild type (Fig 6-A). By contrast, mice infected with AmBRcl.8 (C24SMT) and AmBRcl.6 (C24SMT) showed no macroscopic alterations along the entire experiment and no parasites from these two lines were identified using histology suggesting a fitness loss in these two lines (S11 Fig). Interestingly, material excised from footpad injection sites and associated lymph nodes from all animals was placed into HOMEM yielding proliferative AmBR promastigote cultures from all four AmBR lines and wild type. The AmBRcl.14, AmBRcl.14 PI, AmBRcl.3 and wild type lines reached a density of ∼1×10^5^ cells per mL within 72 hours while AmBRcl.8 and AmBRcl.6 required seven days to reach this density.

While the lower parasite density obtained with AmBRcl.8 and AmBRcl.6 may be explained by the absence of lesion growth observed in mice, the presence of viable parasites recovered from mice tissue with these two lines, which moreover retained their resistant phenotype was unexpected. Previous work with *L. infantum* showed infectious parasites with a unique sterol profiling (129,130,135). Here, it appears that accumulation of ergostanes (96.0 to 96.7%) due to loss of sterol C5 desaturase (C5DS or LOX) is related to the hyper-infectivity observed in AmBRcl.14 and AmBRcl.3. Conversely the accumulation of cholestane type sterols through loss of C24SMT (73.3 to 87.9%) in AmBRcl.8 and AmBcl.6 (Fig 4 & S5-B Table) may also explain the attenuated phenotype observed in these two mutants. Notably, a unique missense heterozygous mutation in both attenuated lines AmBRcl.8 and AmBRcl.6 (S9C Fig) and in AmBRcl.G5 of *L. infantum* (S9A Fig) but not in AmBRcl.14 and AmBRcl.3, was present in the glutathionylspermidine synthase (GSS) gene and in three other genes with unknown function (S10 Fig). GSS is part of the leishmanial multi-protein system (S9B Fig) that protects the parasite against redox species and has been previously linked to virulence in related species (136), however, the impact of these heterozygous changes is not known and other virulence factors cannot be ruled out.

### Retention of resistance *in vivo* and post infection *in vitro* and in macrophages

We also tested whether the parasites from the four lines chosen for *in vivo* infection retained resistance to AmB within the mouse (Fig 6A) and also when recovered from primary lesions (footpad) (Fig 6B & S7 Table) and lymph nodes. After infection, primary lesion size was monitored weekly thereafter and mice were treated with AmB (1 mg per kg) at week eight of infection. In wild type parasites, a plateau in lesion size was attained by fourteen days post-treatment and sustained until the end of the experiment (thirteen weeks). By contrast, mice infected with AmBRcl.14, AmBRcl.3 and AmBRcl.14 PI, the latter recovered from a previous infection, lesion size continued to increase following AmB treatment indicating that these resistant lines were unaffected by this dose of AmB. Due to the absence of increase of lesion in mice infected with AmBRcl.8 and AmBRcl.6 (Fig 6A), retention of resistance in these two lines was assessed *in vitro* as outlined below. Higher dosing was precluded due to the toxicity of the drug, but the data clearly point to all these AmBR parasites being of reduced sensitivity to AmB.

To confirm whether parasites retained resistance after infection, promastigote cultures of the four AmBR lines recovered from footpads and lymph node tissues of infected mice treated with AmB were established and EC_50_ values to AmB measured (S7 Table and S12 Fig). Before infection, AmBR isolates were respectively 11.2-fold (AmBRcl.14, P= 0.0138), 12.3-fold (AmBRcl.3, P= 0.0053), 12.1-fold (AmBRcl.8, P= 0.0062) and 12.8-fold (AmBRcl.6, P= 0.0036) less sensitive to AmB than wild type. Comparable resistance was observed in promastigotes recovered post-infection from all AmBR lines and sub-cultured for two passages (P2) *in vitro* with values of 6.84-fold (AmBRcl.14P2, P= 0.3229), 10.8-fold (AmBRcl.3P2, P= 0.0131), 9.34-fold (AmBRcl.8P2, P= 0.0479) and 12.3-fold (AmBRcl.6P2, P= 0.0093) (Fig 6B). We also tested these four lines that had been recovered from mice (plus the Nys resistant NysRcl.B2 from axenic culture) within infected macrophages and confirmed their resistance phenotype. Interestingly, all lines with mutations in C5DS (and accumulation of ergosta-7,22-dienol) were notably less susceptible (11.20-fold in AmBRcl.14P2, P= 0.0005; 12.2-fold in AmBRcl.3P2, P= 0.0003 and 13.84-fold in Nyscl.B2, P= 0.0001) with respect to their parental wild type (Fig 6C and S7 Table), than either of the two attenuated lines (6.52-fold in AmBRcl.8P2, P= 0.0130 and 4.37-fold in AmBRcl.6P2, P= 0.2100) which had mutations in the C24SMT gene locus (alongside with a deletion of the miltefosine transporter) in which the accumulation of cholesta-5,7,22 was predominant (Fig 4A). Overall, these findings indicate that the retention of the resistant phenotype was stable across all polyene-resistant lines after being recovered from mice (footpads and lymph nodes) (S12 Fig) irrespective of their genomic variants in sterol genes or their virulence.

## Discussion

AmB has emerged as a leading drug to treat leishmaniasis, particularly in its liposomal formulation AmBisome^®^ (8,25,26). Resistance to the drug would impact heavily on efforts to eliminate VL (40, 41). For leishmaniasis, several instances of AmB treatment failure have been reported in the field, involving cases not known to involve immunosuppression by HIV (60, 68) as well as those in HIV infected immunosuppressed patients (47,66,67,137,138).

Moreover, numerous studies have now revealed that resistance can be selected in the laboratory, where the resistance phenotype has frequently been linked to loss of AmB-binding ergostane-type sterols, associated with mutations in enzymes of the sterol pathway. Most often the mutations and loss of expression have arisen in the C24SMT gene locus (60,97,99,139), but in other cases C5DS has been implicated (101) and in one case a loss of C14DM (CYP51) was identified (95). In each case, the mutations lead to loss of ergosterol and an accumulation of other sterol precursors within the pathway, leading to the conclusion that changes in membrane sterols prevent AmB binding to the parasite membrane with the same avidity as to WT cells thus leading to loss of sensitivity. Such mutations have been discovered in laboratory-generated AmB-resistant parasites, as well as clinical isolates of *L. mexicana* (95, 101), *L. donovani* (60,97–99) and *L. infantum* (93).

Apart from our work in which we analysed four AmBR lines (101), previous studies have focused on individual single mutants restricting which restricts the ability to identify different causes of resistance. For this reason, here, we selected a total of 15 new mutants resistant to polyene antimicrobials including 10 *L. mexicana* lines resistant to AmB (AmBR) (7 in HOMEM medium containing FBS and 3 in a serum and lipid free DM). A further four *L. mexicana* lines were selected in HOMEM for resistance to Nys (NysR), a relative of AmB and a single *L. infantum* line was selected for AmBR in HOMEM. Analysis of their respective sterol complements indicated two different profiles had been produced. In both cases, the primary membrane sterol in *Leishmania* spp., ergosterol, was lost and replaced by either ergosta-7, 22-dienol in two of the AmBR *L. mexicana* lines (and in all four NysR lines) selected in HOMEM or cholesta-5,7,22-trienol in five of the HOMEM selected *L. mexicana* AmBR lines and all three of the AmBR lines selected in DM (AmBR-DM), as well as the *L. infantum* AmBR line. In the former group, mutations to C5DS were identified in sequencing the genome while in the latter group mutations to the C24SMT locus were associated. These observations are consistent with previous studies where C24SMT (60,97–99,101) and C5DS (101) have been found associated with development of AmBR. Moreover, experiments in *L. major* where C24SMT (106) and C5DS (107) genes were knocked out of WT parasites also showed that viable parasites with resistance to AmB could be recovered.

Since all of this laboratory-based work has focused on promastigotes forms of the parasites, questions as to their relevance *in vivo* have been made. Purkait et al. isolated parasites from a VL patient who failed treatment and accumulation of sterols indicated changes to C24SMT similar to those found by others in mutants generated under laboratory conditions (97–99, 101). This would indicate the potential of C24SMT mutants to propagate as AmBR cells in patients. *L. major* C24SMT-KO mutants derived by Mukherjee et al. (106) were also of reduced virulence in mice, as were the two lines AmBRcl.8 and AmBRcl.6 with C24SMT mutations we tested here. However, Pountain et al. earlier found a C24SMT mutant that retained virulence in mice. This indicates that it might be mutations other than those to C24SMT *per se* that are responsible for attenuation in mice. Here, for example, we identify changes to the glutathionylspermidine synthase (GSS) gene in both attenuated *L. mexicana* lines (AmBRcl.8 and AmBRcl.6) and in *L. infantum* (AmBRcl.G5) (S9A Fig), although do not yet know whether these changes are associated with attenuation. On the other hand, two separate lines (AmBRcl.14 and AmBRcl.3) with variants in C5DS, both retained virulence and resistance to AmB in mice. In *L. major* C5DS-KO lines displayed only a small loss of virulence in mice (99,106,140), thus C5DS mutations can also be selected, yielding parasites both AmBR and still able to infect mammals. This renders the risk of selection of AmBR in *Leishmania* a genuine risk.

Most studies in AmBR fungi, particularly yeasts including *S. cerevisiae* and *Candida* spp. also showed a tendency for C24SMT and C5DS (ERG6 and ERG3 in fungi, respectively) (75, 77) mutations although changes in other genes of the sterol pathway (ERG-in fungi) such as C14DM (ERG11) (73,77,141,142), C22-sterol desaturase (ERG5) (73, 75) and C-8 sterol isomerase (ERG2) (72, 77) are also involved. In yeast, enhanced oxidative stress resistance has also been implicated in loss of susceptibility to AmB (45,84,85). In *Leishmania*, other studies also pointed to oxidative stress response in AmBR (60,87,91,92,119,134,143). Here we show that the activity of AmB appears to be substantially augmented in rich medium (HOMEM) and associated with enhanced induction of oxidative stress. The activity of AmB against wild type *L. mexicana* grown in HOMEM containing FBS compared with serum-free DM showed the drug to be significantly more potent (3.6 to 4.11-fold) in the rich conditions (Fig 3A,B & D; S3 Table). Metabolomics analysis revealed a significant increase in flux of glucose via the PPP in the cells in rich growth conditions when treated with high concentrations of AmB (Fig 2), which also indicates enhanced oxidative stress causing this increased influx to regenerate NADPH lost in fuelling the redox defence pathways (119–121). Notably, no parasites selected for AmB resistance in DM revealed changes in redox pathway genes, emphasising that environment-specific impacts play key roles in AmB mode of action and resistance.

Overall, this work demonstrates that AmB resistance can be selected relatively easily in *Leishmania* and that changes to sterol profiles predominate. Moreover, we demonstrate that genomic sequencing of multiple lines selected for resistance in parallel can reveal changes that are not identified by studies using single mutants or in which a single gene is knocked out of wild type parasites. As the drug induces oxidative stresses in a manner that is conditional upon the medium in which the parasites grow, then changes to the oxidative stress defence systems can also be important in development of some degree of resistance. When resistance is selected in incremental steps, as done here, a series of events might be behind resistance. In addition to some enzymes involved in the thiol redox systems, three of the lines selected here, as is the case for two earlier demonstrations (101, 102) had lost the miltefosine transporter, which led to cross-resistance to the oral antileishmanial MT in these lines not evident in the other lines. Again, it would seem that the incremental stepwise approach to resistance selection might see some cells changing sensitivity to AmB by other membrane changes associated with loss of the miltefosine transporter, which is a membrane flippase, involved in the architecture of the plasma membrane lipids (93,144–146). The combined roles of redox changes and sterol changes could influence the ability to select and sustain resistance in vivo. For example, two C24SMT mutants were attenuated in mice but an additional mutation in the oxidative defence gene GSS was also found along with a deletion of the MT in each. Further study is required to understand the relative contributions of each. On the other hand, both C5DS mutant lines retained their infectivity in mice. In these, two different independent mutations in the same residue (V74M) and two deletions (M93del and A96del) were found. Whether or not showing virulence, all lines tested in mice retained a stable AmBR phenotype irrespective of their genomic variants in sterol genes or sterol profile. The fact that AmBR has been rare in the field until now might simply be due to the fact that, until recently, it was not widely used, and generally involved protracted dosing regimens.

Extended use and the decision to use a single dose of the drug in monotherapy has great public health benefits (40,41,147,148) but that policy must take into account the fact that selection of resistance is clearly feasible and that resistance can be sustained in parasites that are virulent in mammals. Notably, we observed that, in most cases, AmB resistance is associated with cross-resistance to MT, in some instances, this derived from profound genomic changes such as SNPs and CNVs, as we showed here in three AmBR lines (S7 Fig). Interestingly, here, as in many other studies, AmB resistance is accompanied by increased sensitivity to the antileishmanials PENT and PAR (and to MB). Data from *L. major* evincing a rise in susceptibility to the former were associated with alterations in other membrane lipids (114), which is consistent with interlinked lipid biosynthetic pathways being affected. In the case of PENT, it would seem to be changes in mitochondrial membrane potential due to altered sterol profiles that is responsible (105) and if sterol changes are common then it would be worth considering use of PENT against AmBR lines should they emerge in the field.

## Material and Methods

### Cell culture and polyene-resistant selection

*L. mexicana* (MNYC/BZ/62/M379) and *L. infantum* JPCM5 (MCAN/ES/98/LLM-877) promastigotes were cultured at 25°C and 27°C, respectively, in HOMEM (GE Healthcare and Gibco®) supplemented with 10% v/v heat-inactivated foetal bovine serum (HI-FBS) (Labtech International) or in serum-free defined medium (DM) (149, 150). HOMEM and DM culture media (S2 Table) were supplemented with 100 μg per mL streptomycin, and 100 IU mL−1 penicillin. Parasites were sub-cultured weekly starting at a density of 1 x 10^5^ cells per ml and growth rate, cell density and morphology were determined by light microscopy with a haemocytometer (151, 152). S1 Table shows all the fifteen-individual polyene-resistant lines selected *in vitro* alongside their respective parental wild type, cultured in parallel in the absence of drug. *L. mexicana* cells were selected for resistance by stepwise increasing concentrations of AmB (ten lines) in HOMEM (thirty-two weeks) starting from 50 nM and rising up to 220-300 nM (S1 Fig-A and C) and in DM (fifty weeks) from 100 nM up to 300 nM (S1 Fig-D). Likewise, AmB (one line) for *L. infantum* in HOMEM (fifteen weeks) started at 50 nM up to 100 nM (S1 Fig-E). Nystatin was added to *L. mexicana* (four lines) in HOMEM from 1.5-up to 12.5 µM (S1 Fig-C). Individual clones from each independent resistant line were obtained by limiting dilution assay (LDA). The most resistant clones (n=15) (S1 Table) were selected for further analyses. Retention of resistance (as promastigotes) was confirmed after at least ten passages without drug pressure and post-infection *in vivo* in all resistant lines (n=5) and wild type included in these experiments. Cell lines were preserved in culture medium with 15% DMSO (v/v). Axenic amastigotes were cultured at 32.5 °C with 5% CO_2_ in Schneider’s medium (SDM) pH 5.5, supplemented with HI-FBS (10% v/v), 1.5 mL of hemin (2.5 mg per mL in 50 mM NaOH, 0.003% v/v), 100 μg per mL streptomycin, and 100 IU mL−1 penicillin as described before (153).

### Drug susceptibility assays

Drug susceptibility to various drugs was determined using the Alamar Blue assay (154–156) with some modifications. Briefly, mid-log phase promastigotes (1 x 10^6^ cells per mL) were incubated for 72 hours with different concentrations of drugs serially diluted in a two-fold stepwise fashion in 96-well plates, followed by the addition of resazurin (0.49 mM in 1x phosphate-buffered saline (PBS), pH 7.4) and further incubation for 48 hours. For the experiments of retention of resistance within macrophages, incubation with AmB (AmBisome^®^) and resazurin was for 48 and 6 hours, respectively. Absorbance (fluorescence) was measured using a BMG LabTech Fluostar Optima fluorometer. Intensity was read at λ_EX_ 530 nm and λ_EM_ 590 nm and analysed with Prism 8.0 software to obtain the 50% inhibitory concentration (EC_50_ ± SD, 95% confidence interval) using regression analysis. One-way ANOVA and Tukey’s multiple pairwise comparison tests were performed independently for each compound. All assays were performed in three biological replicates. Unless stated otherwise, all drugs used in these assays were bought from Sigma-Aldrich.

### Infection of mice, macrophages and ethics statement

Eight-week-old female BALB/c mice (from Harlan UK Ltd) were kept at the Central Research Facilities of the University of Glasgow, Glasgow U.K and randomly assigned to either of the treatment groups (wild type versus AmBR lines). Infection of mice was performed by inoculating 2 x 10^6^ stationary phase *L. mexicana* promastigotes resuspended in 100-200 µl of filter sterilised PBS in the left footpad. Mice were treated with six doses of intravenously injected of AmB (1 mg per kg) into the tail vein q.o.d. (every other day) starting at week 8 of infection as described elsewhere (123). Lesion size was assessed weekly before and after chemotherapy (thirteen weeks in total). Amastigotes were recovered post-infection from footpad- and lymph nodes tissue and sub-cultured weekly in Schneider’s medium (SDM) as previously described (153), or transformed into promastigotes and maintained in complete HOMEM. Footpad- and lymph nodes samples were processed for histology and stained with haematoxylin and eosin (H&E) at the Beatson Institute, University of Glasgow, and analysed with light microscopy. Mice were maintained and euthanized following the United Kingdom regulations and the Animals (Scientific Procedures) Act, 1986 (ASPA) under licence PCF371688.

### Retention of resistance to AmB post infection in mice

Retention of resistance to AmB after infection in mice (NysRcl.B2 was from axenic promastigotes culture) was assessed in both axenic promastigotes and in infected macrophages. In the first case, a low passage (P2) of axenic parasites was recovered from mice tissue (as amastigotes) and transformed into promastigotes in HOMEM and drug susceptibility was performed as with previous assays. Retention of AmB resistance within infected macrophages was performed following a method described elsewhere with some modifications (157–159). Briefly, a macrophage-like cell line (RAW 264.7) derived from BALBc/mice was maintained at 37 °C in a humidified atmosphere with 5% CO_2_ in Dulbecco’s Modified Eagle’s Medium (DMEM) with 4.5 g/L glucose, +L-glutamine and 110 mg/L Sodium Pyruvate (Gibco), supplemented with 10% HI-FBS, along with 100 IU and 100 μg/ml each of penicillin and streptomycin. Stationary phase promastigotes (2.5 x 10^6^ per mL) were used to infect adherent macrophage-like cells (2.5 x 10^5^ per mL) at a ratio 10:1 for 24 hours. Infected cells were then washed (3 x) with DMEM to remove extracellular parasites before the addition of variable concentrations of AmB and further incubation for 48 hours. DMEM was then removed and macrophages were lysed using sodium dodecyl sulfate 0.05% (20 µl per well) for 1 min followed by resuspension in complete HOMEM (180 µl per well) and plates were incubated at 26 °C for 48 hours. The presence of promastigotes was assessed with direct microscopy prior to determination of the EC_50_ as described before for drug assays.

### Sterol analysis by GC-MS

GC-MS was performed at Glasgow Polyomics. Sterols were extracted from parasite pellets for Gas chromatography-mass spectrometry (GC-MS) analysis. Mid-log (3 x 10^8^) axenic promastigotes and amastigotes (after transformation in SDM medium as described previously) (153, 160) were washed in PBS and resuspended in 500 ul of KOH-ethanol (20 ml dH_2_O in 30 ml EtOH, 12.5 g KOH) and incubated at 85°C for 1 hour in Pyrex glass tubes. A similar volume of n-heptane was added, and samples were vortexed for 30 seconds. After 20 min the organic layer containing the sterols was transferred into glass borosilicate vials with a Teflon cap (Thermo Fisher®) and stored at −80°C until GC-MS analysis. All samples including blanks (no cells) and pooled samples (a mix with 20 ul from each sample) were in triplicate. Sterols were detected as ester derivatives with trimethyl silane (TMS) and were reported as their underivatized form after normalisation. Briefly, samples were dried with N_2_ flow at 60 °C in glass vials followed by addition of 50 µl of N-methyl-n-trimethylsilyltrifluoroacetamide with 1% 2,2,2-trifluoro-N-methyl-N-(trimethylsilyl)-acetamide, chlorotrimethylsilane (Thermo Scientific), vortexed for 10 secs followed by incubation at 80 °C for 15 mins and allowed to cool down at RT. After this, 50 µl of pyridine and 1 µl of the retention index solution were added and samples were vortexed for 10 secs.

GC was performed in a TraceGOLD TG-5SILMS column with 30 m length, 0.25 mm inner diameter and 0.25 µm film thickness installed in a Trace Ultra gas chromatograph (both from Thermo Scientific) with helium flow rate of 1.0 ml/min, then 1 µl of TMS-derivatised sample was injected into a split/splitless (SSL) injector at 250 °C using a surged splitless injection - splitless time of 30 secs and a surge pressure of 167 kPa. Temperature was increased from 70-up to 250 °C at a ramp rate of 50 °C/min followed by a ramp rate reduction of 10 °C/min reaching a final temperature of 330 °C which was maintained for 3.5 min. Eluting peaks were transferred at an auxiliary transfer temperature of 250 °C to a I0-GC mass spectrometer (Thermo Scientific), with a filament delay of 5 min. Electron ionisation (70 V) was used with an emission current of 50 µA and an ion source that was held at 230 °C. The full scan mass range was 50-700 m/z with an automatic gain control (AGC) of 50%, and maximum ion time of 50 ms. Samples, blanks, pools and standards were loaded into the instrument. Analysis was performed with a TraceFinder v3.3 (Thermo Scientific) and sterols peaks were identified by matching the standards or by comparison to the NIST library.

### Untargeted metabolomics by LC-MS

Mid log *L. mexicana* promastigotes (1 x 10^8^) of all AmB-resistant lines alongside their parental wild-type were treated with AmB (Sigma®) at 5 x their EC_50_ for 15 mins as follows: the amount of AmB for lines selected in HOMEM was 0.3 μM for both parental lines (LmWT1 and LmWT2), 3 μM for AmBRcl.14 and AmBRcl.8, and 1.74 μM for AmBRB2. Both wild type (LmWT-DM) and AmBRA3-DM cultured in DM media were treated with AmB at 1.16 μM and 3.5 μM, respectively.

Metabolite extraction was performed by quenching parasite pellets by rapid cooling at 10°C in dry ice/ethanol bath, culture medium was removed by centrifugation at 1,250 g for 10 minutes at 4°C, then transferred centrifuged twice at 4,500 rpm for 10 minutes at 4°C with a wash in 1ml of cold PBS in between and the pellet was resuspended in 200 ul of monophasic chloroform/methanol/water (CMW 1:3:1) followed by 1 hour shaking (max speed) at 4°C and a final spin at 13,000 rpm for 10 minutes at 4°C. Pooled sample was made by taking 10ul from each sample and blanks were also included. Samples were sealed after adding argon and stored at −80°C until analysis with LC-MS. Identification of metabolites for LC-MS was performed with a ZIC pHILIC column (150 mm × 4.6 mm, 5 μm column, Merck Sequant) coupled to high-resolution Thermo Orbitrap QExactive (Thermo Fisher Scientific) mass spectrometry in both positive and negative ionization modes. Samples were analysed in four replicates and data were processed in Glasgow Polyomics and provided as raw data.

Identification of Liquid Chromatography-Mass Spectrometry (LC-MS) peaks involved mzmatch (161) run though PiMP (118) and IDEOM (162–164) and analysis was conducted in these latter software packages. LC-MS all raw files were submitted to the repository MetaboLights (165) (https://www.ebi.ac.uk/metabolights/) with project number MTBLS2744.

### Genomic DNA extraction and sequencing

Genomic DNA was extracted with the Nucleospin® Tissue kit (Macherey-Nagel) from polyene-resistant and wild-type mid-log promastigotes (1 x 10^7^) washed twice in PBS at 1,250 g for 10 minutes. Library prep and sequencing were performed at Glasgow Polyomics using Illumina sequencer NextSeq 500 yielding 2 x 75 bp paired-end reads. Reference genomes of *L. mexicana* MHOM/GT/2001/U1103 and *L. infantum* JPCM5 (MCAN/ES/98/LLM-877) release 46 were obtained from the TriTrypDB (https://tritrypdb.org). Reads were mapped to their reference genome using BWA-MEM (166) and PCR duplicates were removed using GATK version 4.1.4.1 (167). Variant calling was performed using MuTect2 (168) with the default settings and variant annotation using snpEff (169). Filtered Variant Call Format (VCF) files corresponding to SNPs and indels with an impact on coding sequences were compared between wild type and resistant lines and manually verified using the IGV 2.8.9 visualization tool (170) and eliminated if present in the parental line. Copy ratio alterations were detected using GATK (version 4.2.0.0). Reference genomes were divided into equally sized bins of 1,000 base pairs using PreprocessIntervals tool. Read counts in each bin were collected from alignment data using the CollectReadCounts tool. Copy ratios of a resistant sample over a matched non-resistant sample were obtained from read counts using CreateReadCountPanelOfNormals and DenoiseReadCounts tools. Copy number ratio was obtained by comparing the resistant lines with the corresponding copy number of wild type used as the base line. Genome contigs were segmented using ModelSegments tool from copy ratios of resistant and non-resistant samples. Amplified and deleted segments were identified using CallCopyRatioSegments tool with the default settings. Plots of denoised and segmented copy-ratios were generated using R software. Sequencing raw data was deposited at the NCBI National Center for Biotechnology Information (https://www.ncbi.nlm.nih.gov) with project PRJNA763929 and PRJNA770503.

### Other computational tools

BLAST search was performed against the NCBI database using default settings. ClustalW Omega was used to identify and align sequences. Protein queries-sequences were retrieved from TriTrypDB, Uniprot (https://www.uniprot.org/) and the Protein Data Bank (PDB) (https://www.rcsb.org/). Protein alignment viewers used were Clustal X2 (171). Protein-protein interaction map was created using STRING Database (https://string-db.org) (172–175).

## Acknowledgments

The authors thank Glasgow Polyomics for WGS, GC-MS and LC-MS metabolomics data acquisition.

## Author Contributions

**Conceptualization:** Edubiel A. Alpizar-Sosa <u>orcid.org/0000-0002-0050-6365</u> and Michael P. Barrett.

**Formal analysis:** Edubiel A. Alpizar-Sosa, Nur Raihana Binti Ithnin, Wenbin Wei, Andrew W. Pountain.and Stefan K. Weidt.

**Funding acquisition:** Edubiel A. Alpizar-Sosa, Nur Raihana Binti Ithnin, Paul W. Denny and Michael P. Barrett.

**Investigation:** Edubiel A. Alpizar-Sosa, Nur Raihana Binti Ithnin, Stefan K. Weidt, Anne M. Donachie and Ryan Ritchie.

**Methodology:** Edubiel A. Alpizar-Sosa, Nur Raihana Binti Ithnin, Stefan K. Weidt, Anne M. Donachie and Ryan Ritchie.

**Supervision:** Michael P. Barrett and Richard J. S. Burchmore.

**Writing – original draft:** Edubiel Alpizar and Michael P. Barrett.

**Writing – review & editing:** Edubiel Alpizar, Michael P. Barrett, Emily Dickie and Paul W. Denny.

## Supplementary Information

**S1 Table. List of polyene resistant lines selected for downstream analyses. A)** List of the most resistant clones chosen from each line. **B)** List of individual clones selected for downstream analyses. (XLS).

**S2 Table. Formulation of HOMEM and DM culture media. A)** HOMEM. **B)** Defined Medium (DM). Medium supplemented with serum and antibiotics as described in Material and Methods. (XLS).

**S3 Table. Drugs susceptibility profile (EC50 values) in all polyene resistant lines of *Leishmania* spp.** Mean EC_50_ values (μM) ± Standard Deviation (SD). Except for NMC tested in duplicates, values are from at least three biological replicates. One-way ANOVA was performed independently for each compound to determine differences of the mean between groups. Tukey’s test compared pairwise differences of resistant lines with respect the parental wild type. Statistical difference (P<0.05, 95% Confidence Interval) is shown with stars as follows: ns non-significant or P>0.05; * P≤0.05; ** P≤0.01; *** P≤0.001; **** P≤0.0001.). ND: not determined. AmB (amphotericin B), NMC (natamycin), Nys (nystatin), keto (ketoconazole), MF (miltefosine), PAR (paromomycin), PAT (antimony potassium tartrate), PENT (pentamidine), MB (methylene blue). See Material and Methods for full details. (XLS).

**S4 Table. LC-MS data of AmBR lines cultured in HOMEM and DM.** Metabolites in the pentose-phosphate pathway detected with LC-MS in mid log wild type parasites (1 x 10e8) cultured in HOMEM or DM and treated with high concentrations of AmB for 15 mins (See Material and Methods for a full description). Multivariate data analysis was performed with PiMP analysis pipeline (118) and the Benjamini-Hochberg procedure adjusted raw P-values (q-values) < 0.05 for ANOVA. Each experiment represents one biological replicate (n=4). (XLS).

**S5 Table. Sterol peaks identification and composition (%) found by GC-MS in polyene (AmB and Nys) resistant lines of *Leishmania* spp. A)** Identification of sterol peaks by GC-MS. Standards used were: cholesterol, TMS; desmosterol, TMS; 5α-cholest-7-en-3β-ol, TMS; ergosterol, TMS; stigmasterol, TMS; β-sitosterol, TMS; lanosterol, TMS; FF-MAS (4,4-dimethyl-5α-cholesta-8,14,24-trien-3ß-ol) and zymosterol. FF-MAS: Follicular fluid meiosis-activating sterol and an intermediate in the cholesterol biosynthetic pathway present in all cells. Peaks that did not match any standard were determined by comparing with the NIST spectral libraries with the ion trap mass spectrometer. 5-dehydroepisterol based on literature (de Souza et al. 2009; Xu et al. 2014; Yao et al. 2013; Yao and Wilson 2016). RT: retention time. TMS: trimethylsilyl (for derivatization). **B)** Sterol composition (%) by GC-MS in polyene (AmB and Nys) resistant lines of *Leishmania* spp. promastigotes and amastigotes. Content of each sterol is the percentage of the total of the raw peak area detected per line ± Standard deviation of three independent biological replicates. Sterol content is shown as the total (%) of the raw peak area detected per sample (± SD) from all biological replicates (n=3). (XLS).

**S6 Table. Genotype of mutation identified in sterol genes in polyene resistant and parental wild type of *Leishmania* spp.** LOX (C5DS): Lathosterol oxidase a.k.a. C-5 desaturase (*LmxM.23.1300*); * The genotype in both C24SMT copies is G961/A961 for *LmxM.36.2380* & *LmxM.36.2390* (*LINF_360031200 & LINF_360031300* in *L. infantum*), respectively. ‡ these mutants showed low or no coverage of the intergenic region between both copies of C24SMT (S6 Fig). ND= Non determined due to total absence of coverage at this locus. Changes in the miltefosine transporter (MT) *LmxM.13.1530* are included here and the visualization of genome coverage at this locus is shown in S7 Fig. (XLS).

**S7 Table. Susceptibility to AmB of AmBR *L. mexicana* axenic promastigotes before and after infection in mice and in infected macrophages.** Values in µM, Mean ± Standard Deviation (SD). One-way ANOVA was performed independently for each compound to determine differences of the mean between groups. Tukey’s multiple compared pairwise differences of resistant lines with respect the parental wild type. Statistical difference (P<0.05, 95% Confidence Interval) is shown with stars as follows: ns non-significant or P>0.05; * P≤0.05; ** P≤0.01; *** P≤0.001; **** P≤0.0001. FC: Fold Change of AmBR lines is relative to their respective parental wild type. (XLS).

**S8 Table. Copy Number Variation (CNV) in all polyene resistant lines of *Leishmania* spp.** Copy number ratio was obtained by comparing the resistant lines with the corresponding wild type line which is used as the base line. Gene ID (column A-I) were obtained from TriTrypDB-46. Reference genomes were divided into equally sized bins of 1000 base pairs and the start and end points of the called segments were approximate and not accurate (see Material and Methods for a full description). A visualization of this Table is shown in S8 Fig. (XLS).

**S9 Table. Protein altering mutations (SNPs) identified in all polyene resistant lines of *Leishmania* spp.** Paired end reads were aligned with the corresponding wild type parental line. Reference genome and gene IDs were retrieved from TriTrypDB database (see Material and Methods for a full description). The most relevant mutations are listed in Table 1. (XLS).

**S1 Fig. Selection pressure in polyene-resistant lines of *Leishmania* spp. promastigotes.** *L. mexicana* **(panels A to D)** and *L. infanum* **(panel E)** mid-log promastigotes (5×10^5^) were growth with a stepwise increasing concentration of AmB **(panels A, C-E)** or Nys **(panel B)** added in the cultured medium and indicated with coloured pipelines (left y-axis). The EC_50_ (right-y axis) is shown for wild type (black bars) and resistant lines, AmBR and NysR (coloured bars in all panels). The mean EC_50_ of the parental wild type is shown (horizontal dotted lines). See Material and Methods for a detailed description. (PNG).

**S2 Fig. Growth rate and size of polyene resistant *L. mexicana* promastigotes.** Cell density was measured every 24 hours for 8 days starting from 1×10e5 cells/ml. **A)** AmBR lines in HOMEM. **B)** NysR in HOMEM**. C)** AmBRA4 in HOMEM and AmBRA3-DM in Defined Medium (DM). **D)** Growth kinetics of AmBR lines from panel A (HOMEM) cultured in DM. **E)** Violin plot of the mean size (µm) of four AmBR lines from panel A. The central continuous line within each coloured violin plot is the median of each group. The cell body length was measured from the base of the flagellum until the posterior endpoint of the cell body of promastigotes in the stationary phase. Data were processed with ImageJ software and represent the mean of the sample (n≥ 30). Measurements are the mean (±SD) of three biological replicates. Tukey’s multiple comparison test was used to find pairwise differences between resistant lines and parental wild type. Statistically significant values (P<0.05, 95% Confidence Interval) are indicated with stars as follows: *P ≤ 0.05, **P ≤ 0.01, ***P ≤ 0.001, ****P ≤ 0.0001). (PNG).

**S3 Fig. Fold change in the EC_50_ of antileishmanials in all polyene resistant lines of *Leishmania* spp.** Fold change in EC_50_ is relative to the parental to the respective parental wild type cultured in parallel without drug. Values higher and lower than 1, indicate a decrease and increase in susceptibility, respectively. AmB (amphotericin B),NMC (natamycin), Nys (nystatin), keto (ketoconazole), MF (miltefosine), PAR (paromomycin), PAT (Antimony potassium tartrate), PENT (pentamidine) and MB (Methylene blue). (PDF).

**S4 Fig. Visualization of the genomic region in the sterol gene *LmxM.23.1300* in AmBRcl.3.** The image shows two independent heterozygous mutations, G220A (V74M) and T221A (V74E) in AmBRcl.3. The red square indicates the localisation of both mutation in different reads. Image was produced with WGS data aligned to the reference genome (https://tritrypdb.org) using the IGV_2.8.9 software (http://software.broadinstitute.org/software/igv/). See Fig 5 for a full description of the panels. (PNG).

**S5 Fig. Alignment of the *Leishmania* gene lathosterol oxidase (or sterol C5-desaturase).** *L. mexicana* gene *LmxM.23.1300* (LOX). Black boxes and stars denote His and other residues conserved across species, respectively. Novel variants identified in this study are indicated with black (AmBR lines) and red (NysR line) arrows. Also included (blue box) is the variant from our previous study (Pountain et al. 2019). From top to bottom kinetoplastids listed are *L. mexicana, L. major, L. infantum, L. braziliensis, T. brucei, T. cruzi, Bodo saltans and Crithidia fasciculata*. Also included are *D. rerio* (zebra fish), *M. musculus* (mouse), *Homo sapiens* (human), *S. cerevisiae* (budding yeast), *C. albicans* (pathogenic fungi) and *Arabidopsis thaliana* (plant). Proteins sequences are from TriTrypDB (https://tritrypdb.org) and Uniprot (https://www.uniprot.org/). Alignment was performed using Clustal Ω with default settings (https://www.ebi.ac.uk/Tools/msa/clustalo/). (PNG).

**S6 Fig. Coverage of *LmxM.36.2380* and *LmxM.36.2390* in polyene resistant lines cultured in HOMEM and DM. A-B)** AmBR lines (and their respective parental WT) cultured in HOMEM showing the absence of coverage in this locus in five lines. **C)** AmBR lines and wild type cultured in DM. **D)** Reads coverage in the intergenic region in NysRcl.B2 and its parental WT. **E)** *L. infantum* showing total absence (AmBRcl.G5) and partial coverage (parental WT). Reads were aligned to the reference genome (https://tritrypdb.org) and the image was produced using the software IGV 2.8.9 (http://software.broadinstitute.org/software/igv/). See Fig 5 for a full description of the panels. (PNG).

**S7 Fig. Coverage of the miltefosine transporter of (MT) and adjacent genes in *Leishmania* spp.** The coordinates of a region spanning a genomic region of ∼16 kb and ∼26 kb are shown. Genes are shown at the bottom (blue bars). **A-B)** coverage in NysRcl.B2 and AmBRcl.G5 **C)** In AmBRcl.8 and AmBRcl.6 a total absence of coverage of a region (∼9 kb) comprising *LmxM.13.1530* and adjacent gene downstream *LmxM.13.1540* is shown while small gaps of low coverage are observed downstream from each gene in AmBcl.14, AmBcl.3 and WT. **D)** Absence of a ∼20 kb region comprising four genes (from *LmxM.13.1510* to *LmxM.13.1540*) is shown in AmBRA4. WGS data were aligned to the reference genome (https://tritrypdb.org) and the images were produced with IGV_2.8.9 software (http://software.broadinstitute.org/software/igv/). See Fig 5 for a full description of the panels. (PNG).

**S8 Fig. Copy Number Variation (CNV) in all polyene resistant lines of *Leishmania* spp.** Copy ratio alterations were detected using GATK (version 4.2.0.0). Reference genomes from TriTrypDB46 (https://tritrypdb.org) were divided into equally sized bins of 1000 base pairs (start and end points of the called segments were approximate). CN ratio is obtained by comparing the resistant lines with the corresponding wild type line. Plots of denoised and segmented copy-ratios were generated using R. The thick black lines show the mean copy ratios of corresponding segments. Values of this Fig are provided in S8 Table. (XLS).

**S9 Fig. Visualization of the variants in glutathionylspermidine synthase (GSS). A)** Genomic region showing a heterozygous mutation (boxed) in GSS in AmBRcl.G5 of *L. infantum*. **B)** Protein-protein interaction (PPIs) map of GSS and other enzymes relevant for the trypanothione biosynthesis. GSS (LINF_250031200), GSS: glutathione synthetase (LINF_140015200), TRYS: trypanothione synthetase (LINF_270025600), glutathione peroxidase (LINF_360038100), trypanothione synthetase (LINF_230009800), spermidine synthase (LINF_040010800), trypanosome basal body component (LINF_240028000) and two mannosyltransferases (LINF_360027400 and LINF_120006700). **C)** Genomic region showing the heterozygous mutation in GSS in two lines, AmBRcl.8 and AmBRcl.6 of *L. mexicana*. The reference genome and gene IDs were retrieved from TriTrypDB (https://tritrypdb.org). Images were produced using IGV 2.8.9 (http://software.broadinstitute.org) and STRING (https://string-db.org). See Fig 5 for a full description of the panels A and C. (PNG).

**S10 Fig. Summary of mutations in all polyene resistant lines of *Leishmania* spp. (List of genes).** Gene ID shown on the left were retrieved from TriTrypDB. All polyene resistant lines are shown in the x-axis and the mutation(s) type present in each are coloured coded. This image was produced with R package. See S9 Table for a full list of these variants.

**S11 Fig. Histopathology of primary lesions (footpad) of BALB/c mice inoculated with AmBR lines of *L. mexicana*. A1-2, B1-2)** AmBRcl.14 (yellow) and AmBRcl.3 (orange) infection with intense parasitic load and inflammatory infiltrate within skin histiocytes. **C1-2)** *L. mexicana* wild type (green) causing diffuse inflammatory infiltration. Internalised amastigotes are indicated (black arrows). **D1-2, E1-2)** infection with AmBRcl.8 and AmBRcl.6 (dark and light blue) and discrete inflammatory reaction localised in papillary dermis without visible parasites. Samples are representative of three biological replicates. Hematoxylin-eosin (H&E) stain. Scale bar ∼10 μM Objective ∼63x. (PNG).

**S12 Fig. Retention of resistance (EC50 values) to AmB in *L. mexicana*-AmBR lines recovered from mice tissue (footpad and lymph nodes) after treatment with AmB (1 mg/kg).** Susceptibility to AmB in AmBR *L. mexicana* axenic promastigotes after infection in mice. Parasites were recovered as amastigotes from mice tissue (lymph nodes and footpad) and transformed into promastigotes in HOMEM culture medium. Mice were treated with AmB (1 mg/kg) at week thirteen or left untreated (control group). See Material and Methods for a full description. (PNG).

